# Pantailocins: Phage-derived Bacteriocins from *Pantoea ananatis* and *Pantoea stewartii* subsp. *indologenes*

**DOI:** 10.1101/2023.05.07.539753

**Authors:** Shaun P Stice, Hsiao-Hsuan Jan, Hsiao-Chun Chen, Linda Nwosu, Gi Yoon Shin, Savannah Weaver, Teresa Coutinho, Brian H Kvitko, David A. Baltrus

## Abstract

Phage-derived bacteriocins are highly specific and effective antimicrobial molecules which have successfully been used as prophylactic treatments to prevent phytopathogen infections. Given the specificity of tailocins, a necessary step for broadening the tailocin catalog and for extending applicability across systems and diseases is the screening of new clades of phytopathogens for production of molecules with tailocin-like killing activity. Here, we describe production by and sensitivity of strains to tailocins produced by *Pantoea ananatis* and *Pantoea stewartii* subsp. *indologenes*. Phylogenetic evidence suggests that these tailocins are derived from Myoviridae family phage like many previously described R type syringacins and R type pyocins, but also suggests that cooption from phage occurred independently of previously described tailocins. Since these tailocin encoding loci are present in the same genomic locations across multiple strains of both species and display a level of divergence that is consistent with other shared regions between the genomes and with vertical inheritance of the locus, we refer to them broadly as ‘pantailocins’.

## Introduction

The rise of antibiotic resistance across clinical and agricultural settings as well as increasing appreciation for the importance of limiting off-target effects of broad-spectrum antibiotics on beneficial microbiomes necessitates a search for new, durable, and highly specific antimicrobial compounds(1–3). Phage-derived bacteriocins, referred to as tailocins hereafter, are a class of antibacterial molecules produced by bacterial cells but originally coopted from phage tails(4–6). These molecules are released from cells by lysis to carry out killing in the extracellular environment, maintain high efficiency and specificity in killing of target cells, and have been shown to be effective in preventing infection of plants by phytopathogens(7, 8). Here, we sought to expand the arsenal of known tailocins and identify new types of tailocin molecules that could be used as prophylactic agricultural antibacterials by screening for production and sensitivity across strains of genus *Pantoea* which include several phytopathogens.

Tailocins are antibacterial compounds that closely resemble active phage, except that they largely consist of phage tail proteins (sheath, spike, tail fibers) and therefore lack most if not all proteins involved in capsid production and phage replication(6). Pathways for tailocin production are encoded within the bacterial genome, with an individual cell producing hundreds of these structures before they are released to the environment through cell lysis.

Tailocins are highly efficient in their killing activity, with this lethal activity thought to be due to mechanical disruption of membrane integrity when the spike protein penetrates membranes of sensitive cells(4, 9). Binding of tailocins to target cells has been shown thus far to be primarily controlled by sequence variation in tail fiber proteins and associated chaperones, which interact with sugar moieties in the lipopolysaccharide (LPS) of target cells(10–13). As with all bacteriocins, tailocins are thought to largely target closely related strains to the producer cells, although tailocins from a handful of strains have been shown to have a broader target range (14, 15). Given these properties, tailocins possess incredible potential for development as next generation antimicrobials, with our group and others already demonstrating that tailocins can be applied to plants as a prophylactic treatment to prevent infection by phytopathogens (7, 8).

Much of our knowledge concerning tailocins comes from a handful of *Pseudomonas* species that produce a variety of different tailocin molecules. Specifically, *Pseudomonas aeruginosa* can produce two distinct tailocins, which are referred to as R-type pyocins (which possess a rigid contractile tail) and F-type pyocins (which possess flexible tails)(5, 6). R-type pyocin is considered by many to be the canonical tailocin, is closely related to the Myoviridae phage P2, and has been the focus of research for numerous decades(6, 9, 16). Recently, our group demonstrated that *Pseudomonas syringae* produces an R-type syringacin which we showed to have evolved independently from R-type pyocins, and which is more closely related to the Myoviridae phage Mu than P2(17). Close evolutionary relationships between extant phage and tailocins renders it difficult to definitively identify tailocins by genome sequence alone when evaluating genomic loci in the context of “incomplete” phage regions. Despite such challenges, there have been an increasing number of tailocin producing strains identified, with producers spanning a range of bacterial genera and including both Gram negative and Gram positive cells including multiple species classified as Enterobacterales(18, 19). In one case, *Erwinia* strain Er (now identified as *Pectobacterium carotovorum*) was shown to have the capacity to attach different tail fibers to the tailocin structure, with an invertible region determining which specific tail fiber was encoded by this genome(20).

Given the demonstrated production of tailocins by other Enterobacterales, we sought to screen for tailocin-like killing activity from a variety of *Pantoea* strains isolated from plants.

Here, we report that multiple strains within this genus have the genetic capacity to produce tailocins, and that loci encoding these tailocins maintain at least two distinct killing spectra. Tailocin loci from both species are relatively closely related to each other and likely derived from a common ancestral locus, but are also closely related to extant phage found across Enterobacterales strains and species.

## Materials and Methods

### Screening Pantoea Strains for Tailocin Production

All strains and other genetic materials used in the manuscript are listed in Table 1. We first screened for tailocin-like killing activity against *P. ananatis* strain PNA 97-1R using a previously described overlay assay(21).

**Table 1.**
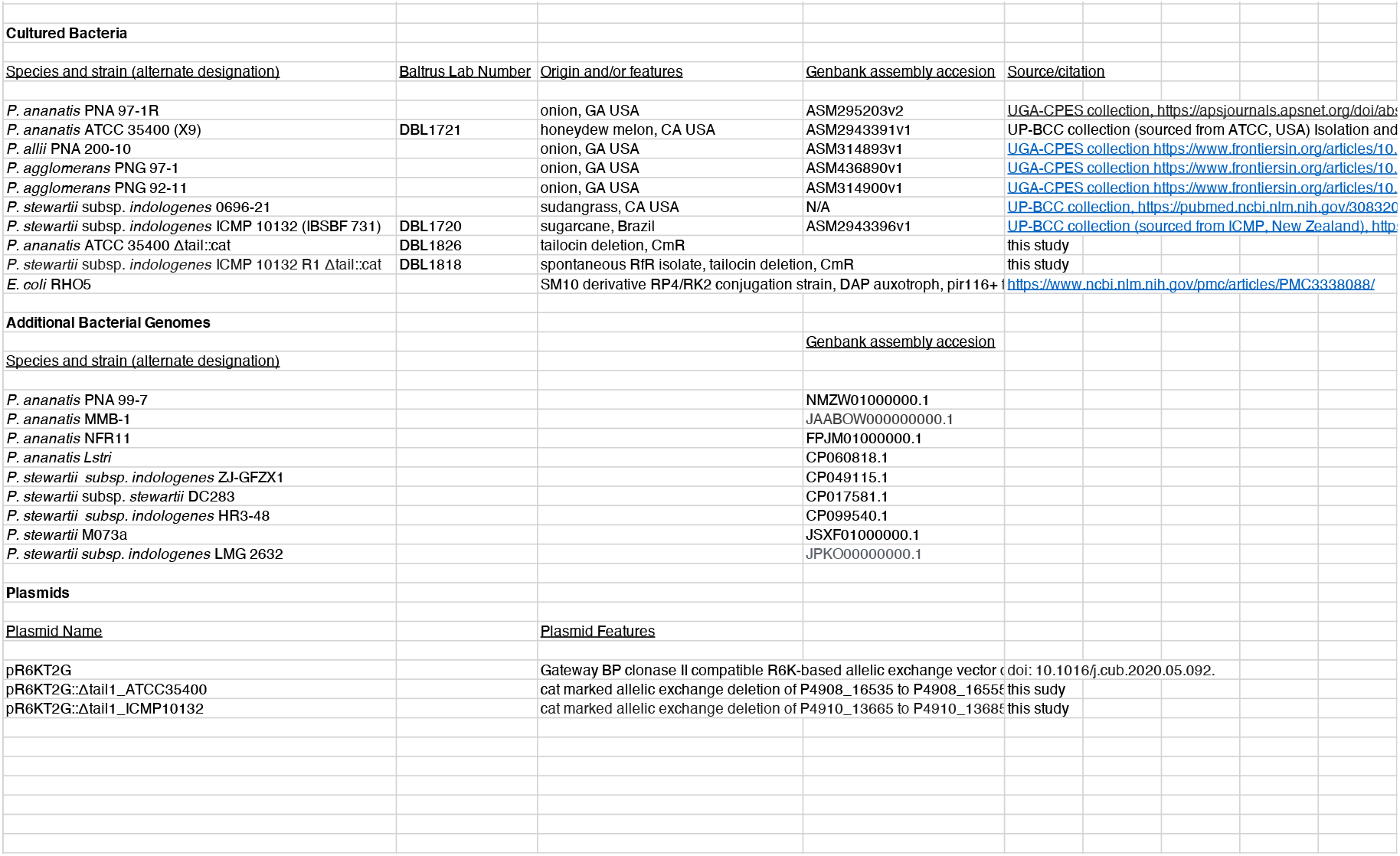
Genetic and Genomic Resources

Supernatants used in this screen were induced for tailocin production and isolated from strains *P. allii* PNA 200-10, *P. agglomerans* PNG 92-11, *P. agglomerans* PNG 97-1, *P. ananatis* ATCC 35400, *P. stewartia* subsp. *indologenes* 0696-21, and *P. stewartii* subsp. *indologenes* ICMP 10132 (which was originally identified as *P. ananatis*, but the isolate within our strain collection has been subsequently identified as a member of *Pantoea stewartii* subsp. *indologenes* based on whole genome sequencing, ANI analysis and subspecies-specific PCR as described in the companion manuscript). We then screened for tailocin production of the same strains against *P. ananatis* strain ATCC 35400. For preparation of tailocins, methods were slightly adapted from previously described reports(21). Briefly, single colonies of each strain were grown overnight in Lysogeny Broth (LB) media at 27°C. Each culture was then back-diluted 1:100 in LB media and grown for 4 hours while shaking at 27°C, at which point Mitomycin C was added to each culture to a final concentration of 0.5 μg/ml. The next day, tailocins were isolated using PEG precipitation and stored for overlay experiments.

For overlay experiments, single colonies of each strain were grown overnight in LB media at 27°C. Each culture was then back-diluted 1:100 in LB media and grown for 4 hours. After 4 hours, 100uL of each culture was added to 3mL molten 1.5% water agar, poured onto LB agar plates, and the top agar was allowed to solidify for ∼15 minutes. At this point, a 5-fold dilution series was prepared for each tailocin in PEG buffer, after which 10μL of the tailocin preparations were added to the overlay plate. Plates were incubated at 27°C for one day at which point plates were observed for tailocin-like killing activity.

### Transmission Electron Microscopy of Potential Tailocins

In all cases, cultures were prepared and PEG precipitated as above to induce and purify tailocins, and then these treatments were visualized using transmission electron microscopy as per (17). Briefly, supernatants from cultures of wild type (DBL1720, DBL1721) and structural mutants (DBL1818, DBL1826) were prepared from strains *P. stewartii* subsp. *indologens* ICMP 10132 and *P. ananatis* ATCC 35400, respectively. Cultures were induced for tailocin production and then concentrated and purified by PEG precipitation. 4 μL of concentrated tailocin samples were placed on glow-discharged carbon-coated grids (Formvar-carbon, 200 mesh copper) for 5 min, then partially blotted to ∼1 μL. Grids were rinsed twice on ddH2O droplet for 5 min. 2% aqueous uranyl acetate was applied on grids for negative staining. Grids were visualized on a JEOL JEM1011 (JEOL USA, Inc, Peabody, MA) transmission electron microscope (TEM) operated at 100kV at the Georgia Electron Microscopy, University of Georgia. Images were taken with a high-contrast 2k x 2k AMT mid-mount digital camera at 50,000× magnification.

### *Genome Sequencing of P. ananatis* ATCC 35400 and *P. stewartii* subsp. *indologens* ICMP 10132

Draft genome sequences were generated for one strain from each *Pantoea* tailocin class, with details of genome sequencing and assembly described in the companion manuscript. For *P. stewartii* subsp. *indologens* ICMP 10132, the Whole Genome Shotgun project has been deposited at DDBJ/ENA/GenBank under the accession JARNMT000000000. The version described in this paper is version JARNMT010000000. For *P. ananatis* ATCC 35400, the Whole Genome Shotgun project has been deposited at DDBJ/ENA/GenBank under the accession JARNMU000000000. The version described in this paper is version JARNMU010000000.

### Identification of Tailocin Locus from Two Different Strains

Each genome sequence was queried for potential phage regions using PHASTER(22), and regions that did not appear to code for complete prophage upon manual inspection were further analyzed. Manual inspection of genomes for both ICMP 10132 and ATCC 35400 highlighted that there exists one potential region in each genome identified as a prophage but which lacks capsid production and phage replication genes and which was therefore consistent with production of a tailocin. Neither of these regions displayed close sequence similarity to previously described tailocins (Figure 5).

### Deletion of Tailocin Structural Loci from Each Strain

For each strain, allelic exchange constructs were designed to disrupt a large portion of the predicted tailocin region by targeting 5 genes predicted to encode tail fibers and baseplate proteins (Figure 2). Allelic exchange in *Pantoea* strains was conducted as described by Stice et al(23). Briefly, 400 bp regions flanking the desired deletion were directly synthesized as dsDNA upstream and downstream of the cat chloramphenicol resistance gene and promoter by Genscript. Synthesized DNA was recombined using BP clonase II into the conditional allelic exchange vector pR6KT2G. Conjugation with the DAP auxotrophic biparental mating strain *E. coli* RHO5 was used to deliver the allelic exchange vectors. Single cross-over merodiploids were recovered by selection on gentamicin and chloramphenicol and double cross-over deletion strains were subsequently recovered by sucrose counter-selection and chloramphenicol selection with screening for loss of the *uidA* color marker on X-gluc and gentamicin sensitivity. Deletions were confirmed by PCR genotyping with independent primers designed to anneal outside the deletion flanks. Tailocin mutant strains were tested for loss of tailocin killing in an overlay assay as well as loss of visible tailocin-like structures by TEM as described above (See supplemental data and figures at doi.org/10.6084/m9.figshare.22596982.v1).

### Phylogenetic Methods

Protein sequences for the baseplate J protein (gp47) and sheath protein from *P. ananatis* ATCC 35400 and *P. stewartii* subsp. *indologenes* ICMP 10132 were used to query the Genbank nr database to find sequences with high similarity from other Enterobacterales species. Protein sequences from these loci were also sampled from multiple *P. ananatis* and *P. stewartii* species that possessed either complete genome sequences or where the *rpoD* to *fadH* locus was contiguous in draft genome sequences. Presumably, homologous sequences were sampled from genomes of strains with previously characterized tailocins and similar phage including *P. aeruginosa* PA01 and P2, *P. syringae* pv. *syringae* B728a and Mu, and from *Pectobacterium carotovorum* strains Er (baseplate) and WPP14 (sheath). Lastly, where identification was possible, J protein and sheath sequences were sampled from additional prophage in *P. ananatis* and *P. stewartii* strains found contiguous with Pantailocin loci in the region between *rpoD* and *fadH*.

Protein sequences were aligned using Clustal Omega with default parameters(24). Alignments were queried in Modeltest to find the most appropriate evolutionary model for inferring maximum likelihood phylogenies(25). RAxML-ng was used to infer phylogenies for each protein alignment, with the LG+G4+F model used for inference for both alignments(26). Bootstrapping was performed for each alignment, and carried out until models converged according to tests within RAxML-ng. Bootstrapping scores were placed on the best tree for each alignment using the Transfer Bootstrap Expectation calculation in RAxML-ng.

## Results

### Identification of Tailocins from Pantoea Strains

Multiple strains classified as Enterobacterales have been shown to possess the ability to produce phage-derived bacteriocins (also known as tailocins), but such molecules have not previously been characterized from *Pantoea* species(18). We find tailocin-like killing activity when supernatant preparations from *P. ananatis* strain ATCC 35400 are tested against *P. ananatis* PNA 97-1R, but not from supernatants from a selection of other strains and species (Figure 1A). This activity forms a crisp border in the overlay assay and does not form single plaques upon dilution, characteristics which are consistent with tailocin-based killing in other systems. We next tested supernatant preparations from each of these strains in overlay assays using ATCC 35400 as the indicator strain. These assays demonstrated that preparations from *P. stewartii* subsp. *indologenes* strains 0696-21 and ICMP 10132 also possess tailocin-like killing activity (Figure 1B) suggesting the presence of a tailocin locus in these strains. Given these results, we independently isolated new tailocin preparations from strains ATCC 35400 and ICMP 10132 and visualized these preparations by transmission electron microscopy. In both cases, visualization of these preparations displayed tailocin-like structures which resemble Myoviridae phage but which lack a capsid (Figures 2A and 2B). Within our preparations, there are also numerous examples of tailocin molecules that have already fired and thus appear as empty sheath molecules. Since we have shown that these tailocins are found in both *Pantoea ananatis* and *Pantoea stewartia*, and that at least a portion of the killing is cross-species, we refer to these tailocins as “Pantailocins”.

**Figure 1.**
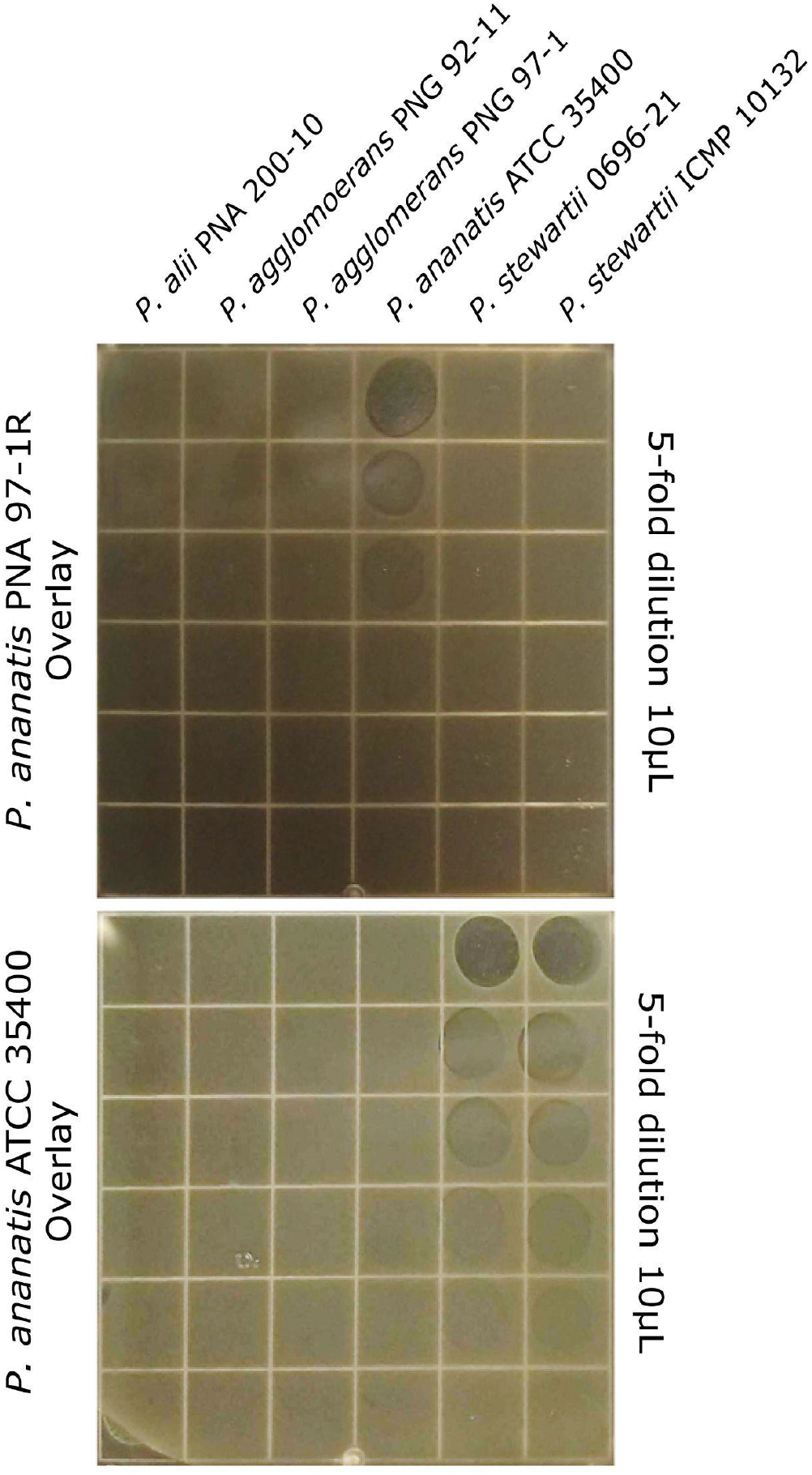
*Pantoea* Strains Display Tailocin-like Killing Activity. A selection of *Pantoea* strains were tested for the ability to produce tailocin-like killing activity against *P. ananatis* PNA 97-1 and *P. ananatis* ATCC 35400. Columns represent antimicrobial activity of preparations from each strain against either of the two target strains, while rows indicate a 5-fold dilution series of the tailocin preparations.

**Figure 2.**
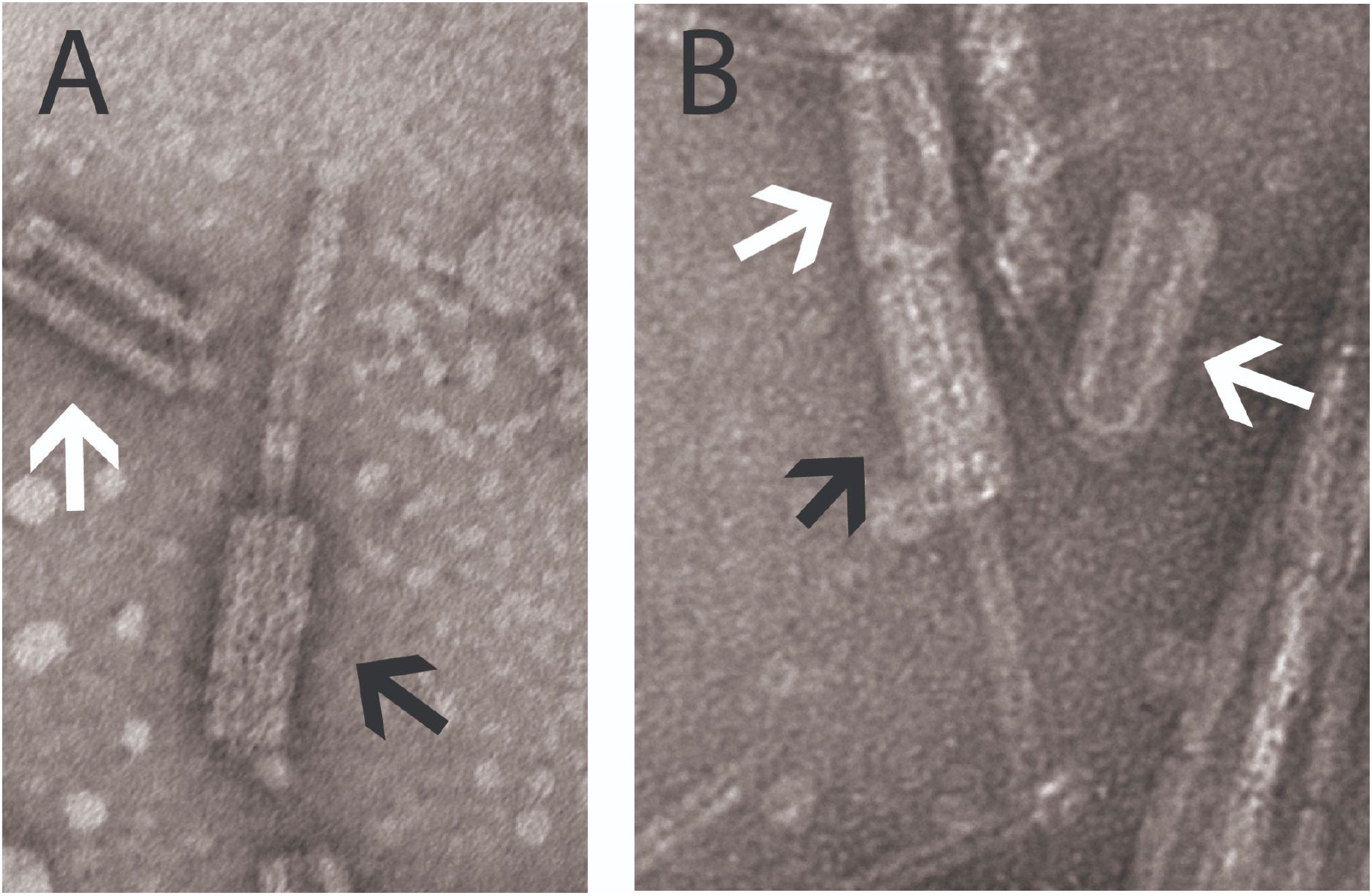
Tailocin-like Molecules Produced by *Pantoea* Strains. We evaluated the production of tailocin molecules by strain A) ATCC 35400 and B) ICMP 10132 using Transmission Electron Microscopy (TEM). Shown are two representative pictures from these TEM preparations. Black arrows indicate tailocins that are ready to fire while white arrows represent tailocin sheaths that have already fired. Original images and additional images can be found at doi.org/10.6084/m9.figshare.22596982.v1

Upon generating draft genome sequences for both ATCC 35400 and ICMP 10132, we searched for potential tailocin encoding loci by first identifying phage regions using PHASTER (22) and manually screening these regions for genomic characteristics consistent with encoding tailocins instead of phage. Predicted tailocins for each strain were found in a region bordered by *dnaG* and *rpoD* as well as a tRNA locus for methionine on one side and by lipoprotein E and *fadH* on the other side. This region in ATCC 35400 appears to contain one predicted tailocin as well as an operon that is predicted to encode proteins involved in nutrient transport. In ICMP 10132, this region appears to encode an additional phage or phage-like structure but lacks genes related to nutrient transport. The conserved border regions (*dnaG/rpoD* and lipoprotein/*fadH*) for the two strains share 80-90% nucleotide similarity, which is roughly the same divergence as most loci predicted to encode tailocins found in both strains (Figure 3). The main difference in alignment of the tailocin regions involves predicted tail fibers, which have been shown to be critical for host targeting specificity of tailocins in other systems. In ATCC 35400, the tail fiber is predicted to be encoded by genes for a receptor binding protein and an associated chaperone.

**Figure 3.**
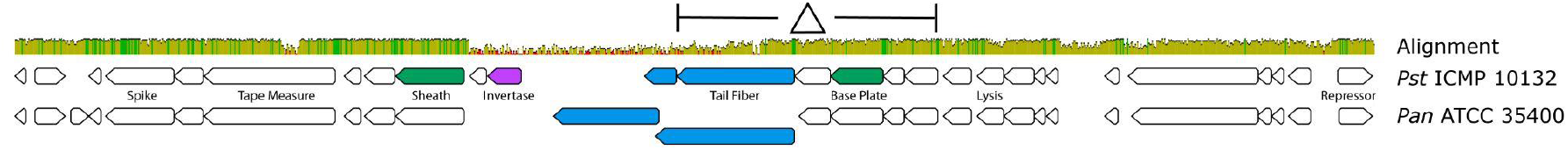
A Schematic of Pantailocin Encoding Loci. Nucleotide sequences from tailocin regions for *Pst*ICMP 10132 and *Pan*ATCC 35400 were aligned against each other, and the alignment and predicted ORFs within each tailocin locus are shown. Nucleotide similarity is displayed by both the height of the alignment similarity bar as well as the colors within the bar itself (green = >80% similarity, yellow = 40-80% similarity, red =<40% similarity). Loci encoding major tailocin genes for both strains are labeled, and the region that was deleted in each strain to create tailocin- mutants is also shown. In both strains the chaperone gene is predicted to be downstream of the tail fiber, but in ICMP 10132 this locus is predicted to be substantially shorter than that found in ATCC 35400. Potential tail fiber proteins and their chaperones are colored blue, the potential invertase is colored purple. Loci used to infer phylogenetic relationships are colored green.

**Figure 4.**
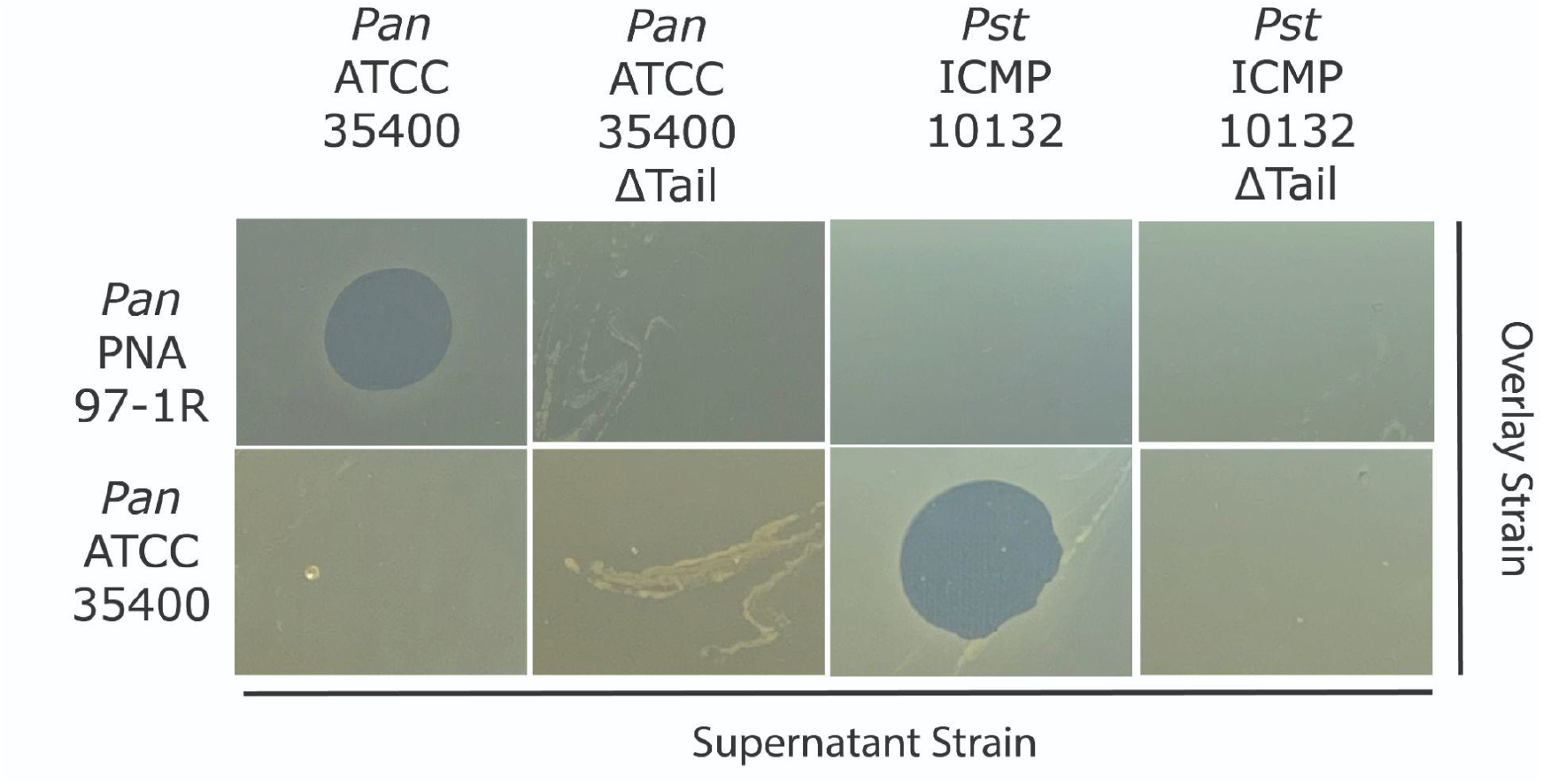
Deletion of Tailocin Structural Genes Abolishes Killing Activity for Both Strains. We created mutants for each strain whereby multiple tailocin structural genes were deleted (see Fig. 3). Tailocin preparations for both wild type and mutant strains were then overlaid onto the two original target strains (PNA 97-1R and ATCC 35400). Tailocin-like activity is demonstrated by clear and crisp killing zones by each strain against its original target, and is abolished in each mutant. Original images can be found at doi.org/10.6084/m9.figshare.22596982.v1.

The tail fiber region in ICMP 10132 is highly diverged from that in ATCC 35400 and stands out from the rest of the predicted tailocin locus because there is little to no nucleotide similarity in these genes between the two strains (Fig. 3). Such a level of divergence is suggestive of localized recombination and horizontal transfer of tail fiber genes for these tailocins as has been demonstrated in *P. syringae* tailocins(12). Strikingly, the recombination point in *Pantoea* appears to be anchored in the N terminus of the tail fiber gene, which was also shown for *P. syringae* tailocins. We also note that in ICMP 10132 there is a gene predicted to encode an invertase immediately adjacent to the Receptor Binding Protein (Rbp) and chaperone. Translation of nucleotide sequences between the invertase and Rbp/chaperone suggests that this locus could encode the C-terminus of an additional Rbp as well as an additional chaperone, which highlights potential for strain ICMP 10132 to alternate tailocin targeting through an invertase based switch.

### Deletions Within Predicted Tailocin Regions Abolishes Killing Activity

To test for associations between these predicted regions and tailocin-like killing activity, we generated multi-gene deletions within predicted tailocin regions for strains ATCC 35400 and ICMP 10132. Specifically, we intended to delete matching stretches of tailocin structural genes in each strain background and we were able to isolate independent strains with syntenic regions deleted in both ATCC 35400 and ICMP 10132. In each case, deletions eliminated tailocin-like killing activity present in the wild type strains when evaluated using overlay assays (Figure 5). We have also visualized preparations from these deletion strains using TEM and as part of the same preparations used to visualize the wild type strains as above, and note that there are few (if any) tailocin-like structures created in these deletion mutants (See supplemental data and figures at doi.org/10.6084/m9.figshare.22596982.v1).

**Figure 5.**
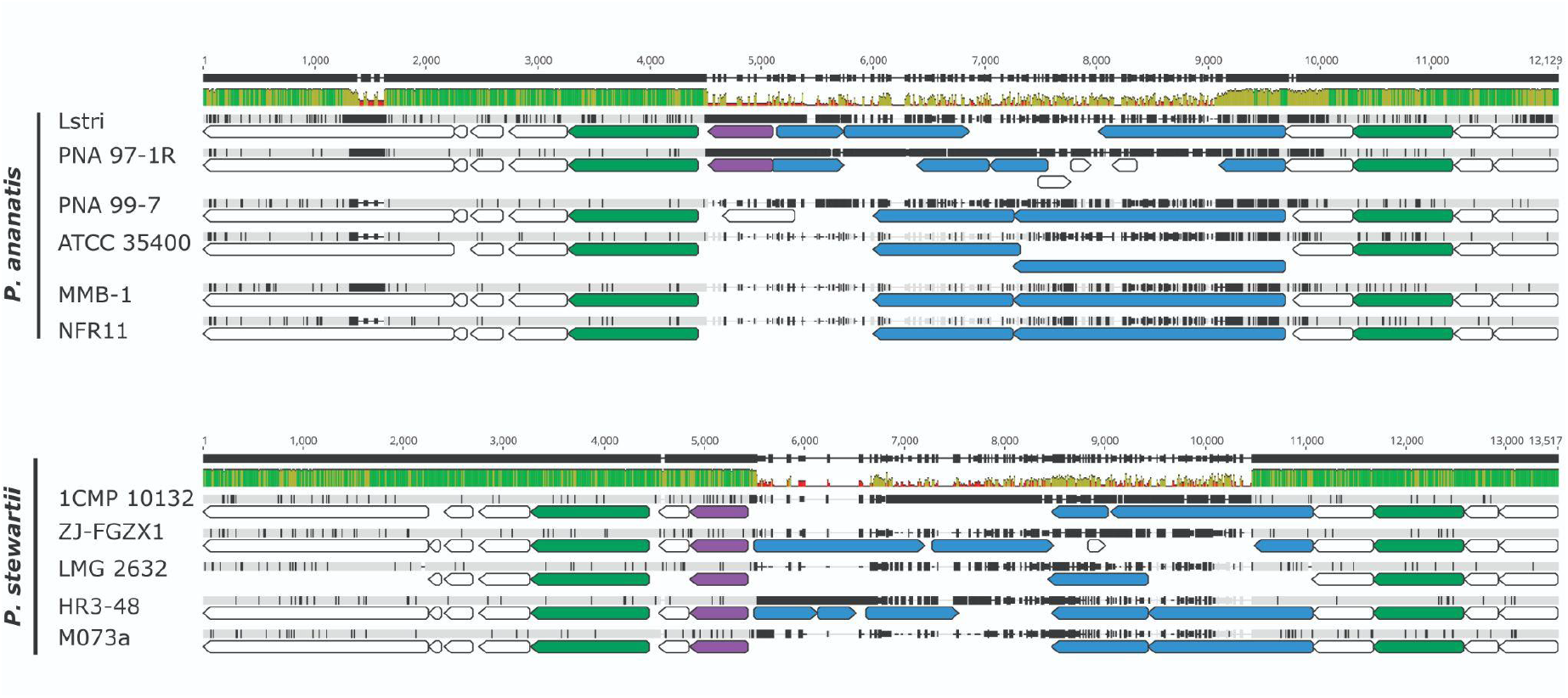
Alignment of Pantailocin Loci Across *P. ananatis* and *P. stewartii* Strains. Nucleotide sequences for regions spanning the tape measure protein to baseplate protein V from predicted tailocin regions for numerous *P. ananatis* and *P. stewartii* strains were aligned, and the alignment and predicted ORFs within each tailocin locus are shown. Nucleotide similarity is displayed by both the height of the alignment similarity bar as well as the colors within the bar itself (green = >80% similarity, yellow = 40-80% similarity, red =<40% similarity). Potential tail fiber proteins and their chaperones are colored blue, the potential invertase is colored purple. Loci used to infer phylogenetic relationships are colored green. Original alignments and phylogenetic data can be found at doi.org/10.6084/m9.figshare.22596982.v1

### Tailocin Loci are Predicted to be Found in Additional P. ananatis and P. stewartii Strains

To gauge the distribution of potential tailocin loci across both *P. ananatis* and *P. stewartii*, we investigated the region in between *rpoD* and *fadH* in closed genomes for multiple strains of each species (Table 1) with a focus on strains with complete genome sequences where the entirety of this region was contiguous. In many strains, we found evidence for independent prophage integration at this site, similar to what is found in ICMP 10132, as well as a relatively high level of presence/absence diversity in genes between *rpoD* and the predicted start of the tailocin loci. However, we did find evidence for pantailocin encoding loci similar to those described for ATCC 35400 and ICMP 10132 in this region of every genome of *P. ananatis* investigated and in a majority of genomes classified as *P. stewartii* subsp. *indologenes*. The only analyzed *P. stewartii* genome that appeared to lack a tailocin locus between *rpoD* and *fadH* was DC283, which is a reference strain used to study virulence in *P. stewartii* subsp. *stewartii*. As a further comparison of tailocin loci, we aligned regions spanning from the tailocin tape measure protein through baseplate protein V (Fig. 5). As shown in Fig. 3 for ATCC 35400 and ICMP 10132, tailocin regions align quite well across *P. ananatis* and *P. stewartii* strains except for extensive diversity in the loci coding for predicted receptor binding proteins (Fig. 5). Moreover, many of the examined tailocin loci across *P. ananatis* strains and all *P. stewartii* strains including the confirmed ICMP 10132 pantailocin, appear to encode an invertase that presumably enables inversion within the tailocin region. Such a molecular switch could impart in the genetic capability for strains containing this invertase to encode multiple tail fibers with different specificities.

### Pantoea Tailocin Loci are Divergent from Previously Described Tailocins

To better understand the evolutionary relationships between tailocin loci encoded by *Pantoea* and those described from other systems, we inferred phylogenies for predicted baseplate protein (J-like, gp47) alleles as well as the predicted sheath proteins encoded by these systems, by related phage, and by other described tailocin systems. Phylogenies for both proteins show consistent relationships between alleles of the proteins investigated with strong bootstrap support for many of the major clades and divisions. The tailocin baseplate and sheath proteins from *Pantoea* strains ATCC 35400 and ICMP 10132 form clades with predicted Pantailocin loci from closely related strains within the same species. Together, loci from *Pantoea* tailocins form a clade to the exclusion of proteins from other tailocins: including the R pyocin from *P. aeruginosa*, R syringocin from *P. syringae*, and the previously described *Pectobacterium* R carotovoricin. Our phylogenies also reflect the idea that the R pyocin from *P. aeruginosa* is closely related to loci from phage P2 and that the R syringocin is closely related to loci from phage Mu(16, 17). However, the proteins most closely related to those found in the tailocins of *Pantoea* are from regions of other Enterobacterial strains that appear to encode extant prophage. Specifically, manual inspection of these regions from additional Enterobacterial strains shows that the baseplate/sheath loci from these genomes are colocalized with numerous genes predicted to encode a capsid. Moreover, loci from *Pantoea* tailocins are more closely related to those from *Erwinia* phage ENT90 than to P2 (again to the exclusion of the *P. aeruginosa* R pyocin proteins).

Previously described tailocin loci from *Pseudomonas aeruginosa* and *Pseudomonas syringae* have independently arisen through cooption of different progenitor phage from the Myoviridae family. In the case of *P. aeruginosa*, the most well described closely related phage to R-type pyocins is P2 (16) while the most well described closely related phage to R-type syringacins is Mu(17). Phylogenetic relationships suggest that, like the R-type pyocins of *P*.*aeruginosa*, tailocins from *Pantoea* are derived from a phage that resembles P2 rather than Mu but also that this may also represent an independent cooption of phage into tailocins given close relationships with extant phage to the exclusion of proteins from phage P2 as well as the R-type pyocins.

However, definitive characterization of potential origins of pantailocins will require additional sampling and evolutionary analyses as it is also possible that recombination between the *Pantoea* tailocins and extant phage is obscuring definitive evolutionary signals. We also note that even though a tailocin containing an invertase switch has previously been shown for carotovoricin Er (20), a tailocin encoded by *Pectobacterium carotovorum*, the predicted tailocin in ICMP 10132 is highly divergent to the previously described locus from *Erwinia* strain Er (Fig. 6) and that the *Pectobacterium* tailocin is more close in sequence to *P. aeruginosa* R-type pyocins than the *Pantoea* tailocins described here. Therefore, tailocins from *Pantoea* and from *Pectobacterium* and their invertible tail fibers also appear to have independently arisen from divergent ancestor phage.

**Figure 6.**
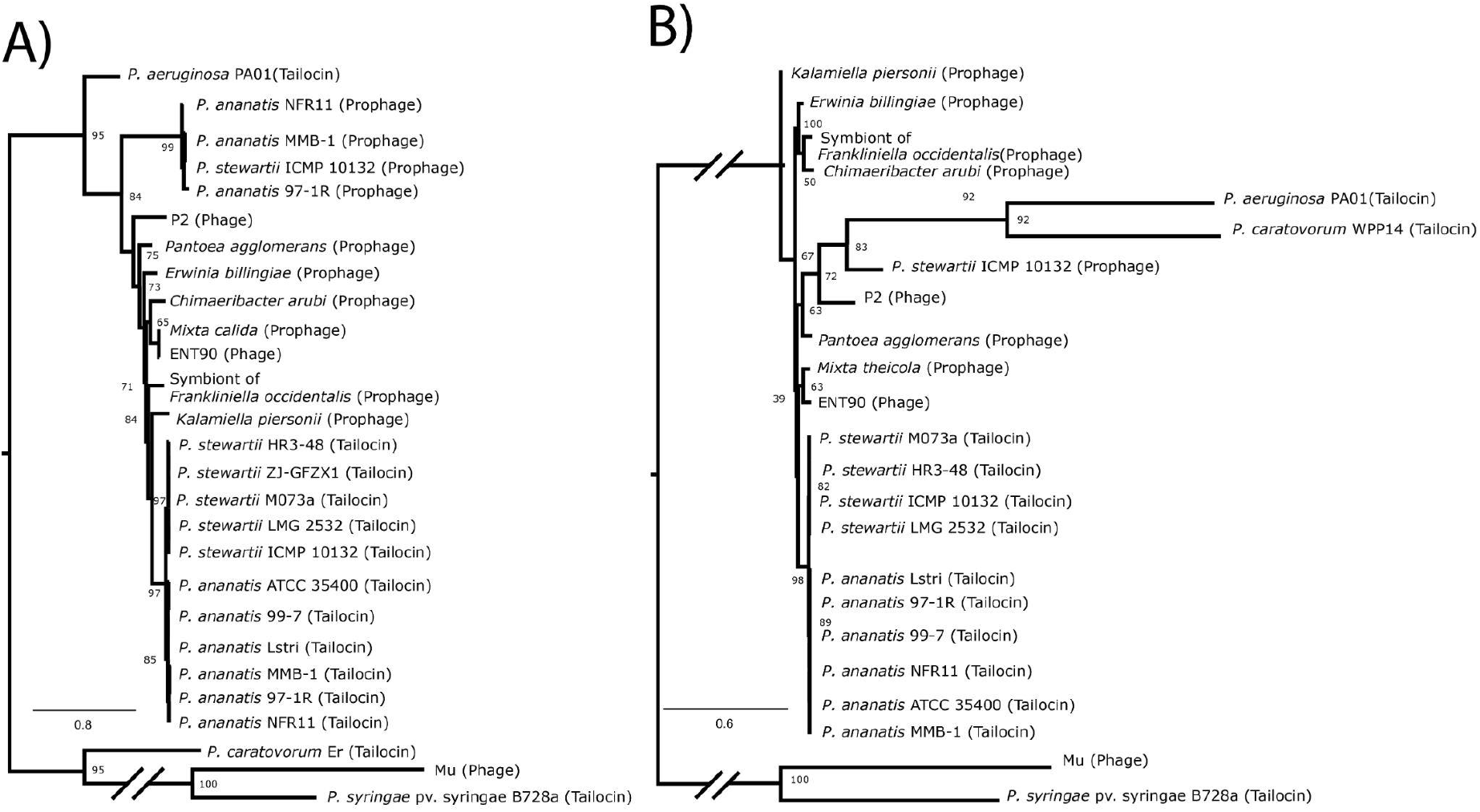
Phylogenetic Relationships of Pantailocin Loci Compared to Other Phage and Tailocins. Maximum likelihood phylogenies for the A) J plate and B) Sheath protein were inferred using alignments from predicted and known tailocins as well as related phage and prophage. Bootstrap values are shown for major nodes, but where not shown bootstrap values are generally <75 (e.g. in the *P. ananatis* and *P. stewartii* tailocin clades where there is little resolution to determine phylogenetic relationships).

### Conclusions

We have described new tailocin loci found within the genomes for strains characterized as both *Pantoea ananatis* and *Pantoea stewartii* subsp. *indologenes*. These tailocins are encoded by a locus found within the same approximate genomic context across strains (between *rpoD* and *fadH* and adjacent to a gene for tRNA_Met_), and share a level of divergence similar to other shared genes throughout the genome (∼85-90%). Identification of these loci opens up the possibility that these tailocin molecules could be developed as new prophylactic treatments to prevent infection of crops by *Pantoea* or potentially as a means to limit spread of these strains by various vectors.

## Notes

### Competing Interest Statement

The authors have declared no competing interest.

## References

1. Manesh A, Varghese GM, CENDRIC Investigators and Collaborators. 2021. Rising antimicrobial resistance: an evolving epidemic in a pandemic. Lancet Microbe 2:e419–e420.

2. Chinemerem Nwobodo D, Ugwu MC, Oliseloke Anie C, Al-Ouqaili MTS, Chinedu Ikem J, Victor Chigozie U, Saki M. 2022. Antibiotic resistance: The challenges and some emerging strategies for tackling a global menace. J Clin Lab Anal 36:e24655.

3. Uddin TM, Chakraborty AJ, Khusro A, Zidan BRM, Mitra S, Emran TB, Dhama K, Ripon MKH, Gajdács M, Sahibzada MUK, Hossain MJ, Koirala N. 2021. Antibiotic resistance in microbes: History, mechanisms, therapeutic strategies and future prospects. J Infect Public Health 14:1750–1766.

4. Taylor NMI, van Raaij MJ, Leiman PG. 2018. Contractile injection systems of bacteriophages and related systems. Mol Microbiol 108:6–15.

5. Ghequire MGK, De Mot R. 2015. The Tailocin Tale: Peeling off Phage Tails. Trends Microbiol 23:587–590.

6. Scholl D. 2017. Phage Tail-Like Bacteriocins. Annu Rev Virol 4:453–467.

7. Baltrus DA, Clark M, Hockett KL, Mollico M, Smith C, Weaver S. 2022. Prophylactic Application of Tailocins Prevents Infection by Pseudomonas syringae. Phytopathology 112:561–566.

8. Príncipe A, Fernandez M, Torasso M, Godino A, Fischer S. 2018. Effectiveness of tailocins produced by Pseudomonas fluorescens SF4c in controlling the bacterial-spot disease in tomatoes caused by Xanthomonas vesicatoria. Microbiol Res 212–213:94–102.

9. Ge P, Scholl D, Prokhorov NS, Avaylon J, Shneider MM, Browning C, Buth SA, Plattner M, Chakraborty U, Ding K, Leiman PG, Miller JF, Zhou ZH. 2020. Action of a minimal contractile bactericidal nanomachine. Nature 580:658–662.

10. Heiman CM, Maurhofer M, Calderon S, Dupasquier M, Marquis J, Keel C, Vacheron J. 2022. Pivotal role of O-antigenic polysaccharide display in the sensitivity against phage tail-like particles in environmental Pseudomonas kin competition. ISME J 16:1683–1693.

11. Köhler T, Donner V, van Delden C. 2010. Lipopolysaccharide as shield and receptor for Rpyocin-mediated killing in Pseudomonas aeruginosa. J Bacteriol 192:1921–1928.

12. Baltrus DA, Clark M, Smith C, Hockett KL. 2019. Localized recombination drives diversification of killing spectra for phage-derived syringacins. ISME J 13:237–249.

13. Jayaraman J, Jones WT, Harvey D, Hemara LM, McCann HC, Yoon M, Warring SL, Fineran PC, Mesarich CH, Templeton MD. 2020. Variation at the common polysaccharide antigen locus drives lipopolysaccharide diversity within the Pseudomonas syringae species complex. Environ Microbiol 22:5356–5372.

14. Weaver SL, Zhu L, Ravishankar S, Clark M, Baltrus DA. 2022. Interspecies Killing Activity of Pseudomonas syringae Tailocins. bioRxiv.

15. Yao GW, Duarte I, Le TT, Carmody L, LiPuma JJ, Young R, Gonzalez CF. 2017. A Broad-Host-Range Tailocin from Burkholderia cenocepacia. Appl Environ Microbiol 83.

16. Nakayama K, Takashima K, Ishihara H, Shinomiya T, Kageyama M, Kanaya S, Ohnishi M, Murata T, Mori H, Hayashi T. 2000. The R-type pyocin of Pseudomonas aeruginosa is related to P2 phage, and the F-type is related to lambda phage. Mol Microbiol 38:213–231.

17. Hockett KL, Renner T, Baltrus DA. 2015. Independent Co-Option of a Tailed Bacteriophage into a Killing Complex in Pseudomonas. MBio 6:e00452.

18. Patz S, Becker Y, Richert-Pöggeler KR, Berger B, Ruppel S, Huson DH, Becker M. 2019. Phage tail-like particles are versatile bacterial nanomachines - A mini-review. J Advert Res 19:75–84.

19. Gebhart D, Williams SR, Bishop-Lilly KA, Govoni GR, Willner KM, Butani A, Sozhamannan S, Martin D, Fortier L-C, Scholl D. 2012. Novel high-molecular-weight, Rtype bacteriocins of Clostridium difficile. J Bacteriol 194:6240–6247.

20. Nguyen HA, Tomita T, Hirota M, Kaneko J, Hayashi T, Kamio Y. 2001. DNA inversion in the tail fiber gene alters the host range specificity of carotovoricin Er, a phage-tail-like bacteriocin of phytopathogenic Erwinia carotovora subsp. carotovora Er. J Bacteriol 183:6274–6281.

21. Hockett KL, Baltrus DA. 2017. Use of the Soft-agar Overlay Technique to Screen for Bacterially Produced Inhibitory Compounds. J Vis Exp https://doi.org/10.3791/55064.

22. Arndt D, Grant JR, Marcu A, Sajed T, Pon A, Liang Y, Wishart DS. 2016. PHASTER: a better, faster version of the PHAST phage search tool. Nucleic Acids Res 44:W16–21.

23. Stice SP, Thao KK, Khang CH, Baltrus DA, Dutta B, Kvitko BH. 2020. Thiosulfinate tolerance is a virulence strategy of an atypical bacterial pathogen of onion. Curr Biol 30:3130-3140.e6.

24. Sievers F, Higgins DG. 2018. Clustal Omega for making accurate alignments of many protein sequences. Protein Sci 27:135–145.

25. Darriba D, Posada D, Kozlov AM, Stamatakis A, Morel B, Flouri T. 2020. ModelTest-NG: A new and scalable tool for the selection of DNA and protein evolutionary models. Mol Biol Evol 37:291–294.

26. Kozlov AM, Darriba D, Flouri T, Morel B, Stamatakis A. 2019. RAxML-NG: a fast, scalable and user-friendly tool for maximum likelihood phylogenetic inference. Bioinformatics 35:4453–4455.

